# Multi-Omics Analysis of Heat Stress-Induced Memory in Arabidopsis

**DOI:** 10.1101/2025.06.19.660594

**Authors:** Venkatesh P. Thirumalaikumar, Liang Yu, Devender Arora, Umrah Mubeen, Andrew Wiszniewski, Dirk Walther, Patrick Giavalisco, Saleh Alseekh, Andrew DL Nelson, Aleksandra Skirycz, Salma Balazadeh

## Abstract

In their natural environment, plants experience temperature fluctuations, including intensified heat waves driven by climate change, which pose significant threats to their productivity. To adapt, plants have evolved diverse mechanisms to withstand heat stress (HS), minimizing potential damage and ensuring survival. One such adaptation is acquired thermotolerance (AT), where prior exposure to HS primes plants to withstand subsequent severe HS. AT can persist for several days and involves a recovery period during which plants establish heat stress memory (HSM), reorganize cellular processes, and strengthen stress resilience. The molecular mechanisms underlying HSM remain the subject of active investigation. In this study, we employ a high-throughput comparative multi-omics approach to unravel the transcriptome, metabolome, and proteome of *Arabidopsis thaliana* seedlings during distinct intervals of the HSM phase. Our findings provide insights into the intricacies of HS recovery and the memory process. Notably, distinct temporal responses emerge at both the transcriptional and protein levels during the early and late recovery phases. Transcripts associated with HSM are upregulated during the early HS recovery phase, indicating a rapid response crucial for initial memory formation, while corresponding protein levels remain elevated throughout the recovery period, supporting memory consolidation. Additionally, metabolite profiles reveal distinctive patterns across the HS memory phase. This marks the first detailed multi-omic analysis of the HSM phase in Arabidopsis seedlings, providing insights into the multifaceted nature of this complex process. These comprehensive datasets hold promise in elucidating regulators of HS resilience, thereby enhancing efforts in breeding HS-tolerant crops

## Background and Summary

As sessile organisms, plants have developed numerous mechanisms to perceive and respond to fluctuating environmental conditions. In their natural habitats, stressors such as drought, temperature extremes, and salinity frequently arise and recur throughout their life cycle (Bäurle, 2016; Hilker et al., 2016; Balazadeh, 2022; Staacke et al., 2025). To cope with such stressful conditions, plants develop acquired tolerance through priming, where prior stress exposure enhances their ability to mount a more efficient response to subsequent stress events. Priming triggers a new cellular homeostasis state that can persist during the recovery period, even after the priming stimulus is removed. This state enhances the plant’s preparedness for future stress events, a phenomenon known as memory (Bäurle, 2016). Evidence shows that stress-priming, followed by the establishment of memory is beneficial for stress survival in plants, particularly in the short term (Stief et al., 2014, Brzezinka et al., 2016, Feng et al. 2016; Sedaghatmehr et al., 2016,2019, 2021; Ganguly et al. 2017).

High temperatures adversely affect plants’ growth and productivity (Balazadeh, 2022). Plant cells redirect their metabolism and resources to mitigate or prevent the damages caused by heat stress (HS) (Mittler et al., 2012). Prolonged exposure to HS disrupts proteome integrity through protein misfolding and/or denaturation, leading to cell toxicity and death (Chen et al., 2011; Zhou et al. 2014a,b). To cope with HS, plants activate their heat stress response (HSR) pathways such as (1) increasing their antioxidant activities, (2) activating and/or repressing HS-related transcription factors (TFs) in conjunction with the accumulation of heat shock proteins (HSPs), (3) adjusting their chromatin architecture through activation or repression of the epigenetic regulators (Mittler et al., 2012, Bäurle, 2016; Zhou et al. 2014a,b), among other strategies.

HSPs are grouped into different families, considering their molecular masses: HSP70, HSP90, HSP100/ClpB (Hsp101), and small HSP (sHSP) families. Small HSPs with monomeric molecular mass ranging from 12 to 42□kDa, belong to an evolutionarily conserved family characterized by a common α-crystalline domain located at the C-terminal part (Sun and MacRae, 2005). Some of these sHSPs are targeted to different cellular compartments, including chloroplasts, mitochondria, cytoplasm and the endoplasmic reticulum (ER), with several studies confirming their role in protecting organelle-specific proteins during or after HS (Perez et al. 2009; Escobar et al. 2021; Huang et al. 2024).

Cellular and molecular connections related to plant growth and stress defense are well described using biological networks. Often, these biological networks are constructed based on one or two omics strategies, which may not provide a holistic view of the responses in context (Clark et al., 2023). It is increasingly recognized that a better understanding of the biological processes can be achieved through multi-omics approaches. Multi-omics studies offer a more comprehensive platform for deciphering the mechanisms and pathways involved in various biological models (Clark et al. 2023; Gupta et al. 2024).

While numerous studies have described the HS responses during or shortly after HS in Arabidopsis, there has been a lack of integrated multi-omics research focusing on events during the HS recovery period and the establishment of memory. Here, we present the first detailed multi-omic analysis of the HSM phase in Arabidopsis seedlings, providing insights into the multifaceted nature of this complex process.

Figure 1 shows the experimental conditions and pipeline employed for various omics analyses (transcriptome, proteome, and metabolite profiling). We use the term HSM phase to describe the period during which a plant recovers from HS priming while simultaneously consolidating and retaining the memory of that HS. We employed untargeted proteomics to identify differentially expressed proteins and those that potentially regulate HSM in Arabidopsis by comparing the proteomes of primed and control plants. In our experiment, we detected ∼5200 proteins with 1%FDR. The quality of the treatment on the proteomic datasets was evaluated by principal component analysis (PCA) (**Figure 2A, C, E, G**), and the differentially regulated proteins were visualized by volcano plots (**Figure 2B, D, F, H**). As expected, proteomics data showed HSPs (such as Heat shock protein 21 (HSP21), Heat shock protein 18 (HSP18.1), Heat shock protein 15.7 (HSP15.7), Heat shock protein 90.1 (HSP90.1), Heat shock protein 22 (HSP22) and other stress response proteins such as Frigida-essential 1 (Fes1A) are induced and remain upregulated at the early phases of HSM phase (up to 2 day at HSM phase). This effect becomes subtler during the later stages of the HSM phase (2–3 days after the priming stimulus). We also identified a set of proteins that are induced and remain elevated during the early stages of HSM, which we refer to as HSM-associated proteins. Notable examples include HSP21, HSP15, ClPB1, and FES1A. Interestingly, the abundance of most HSM-related proteins appeared to decrease or become less pronounced in the later stages of the HSM phase, indicating the reset of the memory phase. Notably, their corresponding transcripts were upregulated only in the early phases of HSM. The transcriptomic analysis is presented in **Figure 3**, along with a principal component analysis score plot that provides and overview of transcriptome-based sample similarities and differences (**Figure 3A**).

**Figure 1.**
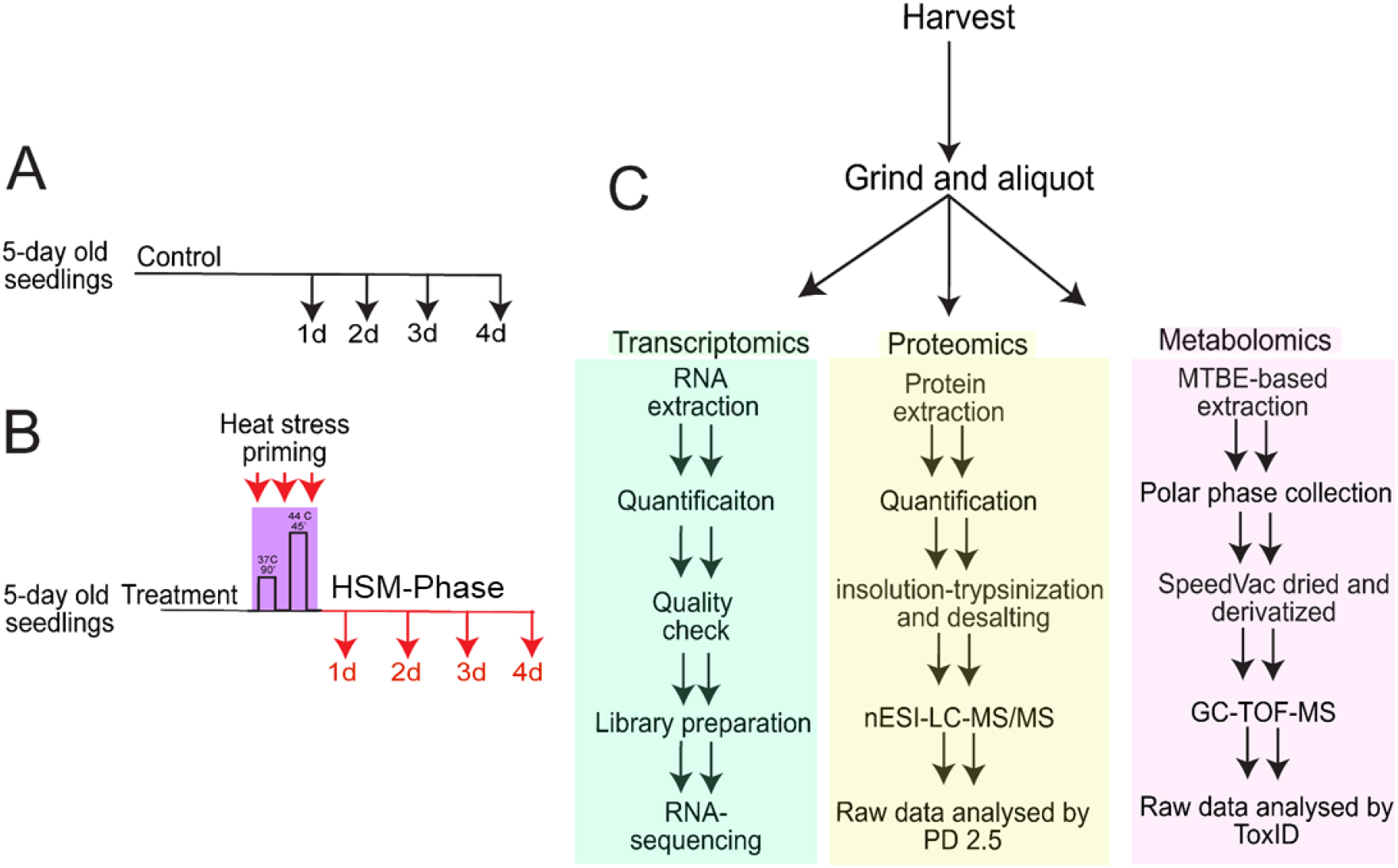
Graphical illustration of the experimental setup and sample collection. (**A)** Five-day-old *Arabidopsis thaliana* seedlings were subjected to a two-step HS priming regime of 37^°^C for 90 min, followed by a room temperature recovery for 90 min and then 44^°^C for 45 min. After the priming stimuli, the seedlings were returned to regular growth conditions. The seedlings were harvested at different time points as plants entered the HS recovery/memory phase, as indicated. (**B**) Seedlings not subjected to HS priming treatment but experienced the same light regime changes were harvested as control seedlings at the indicated t time points. (**C**) Schematic representation of the experimental pipeline. Seedlings were ground using liquid nitrogen conditions and aliquoted for the RNA extraction proteomic and metabolomic analysis. For proteomics and metabolomics MTBE-methanol method (REF) was used. Polar fractions were analyzed using GC-MS, while protein fractions were analyzed using LC-MS/MS.

**Figure 2.**
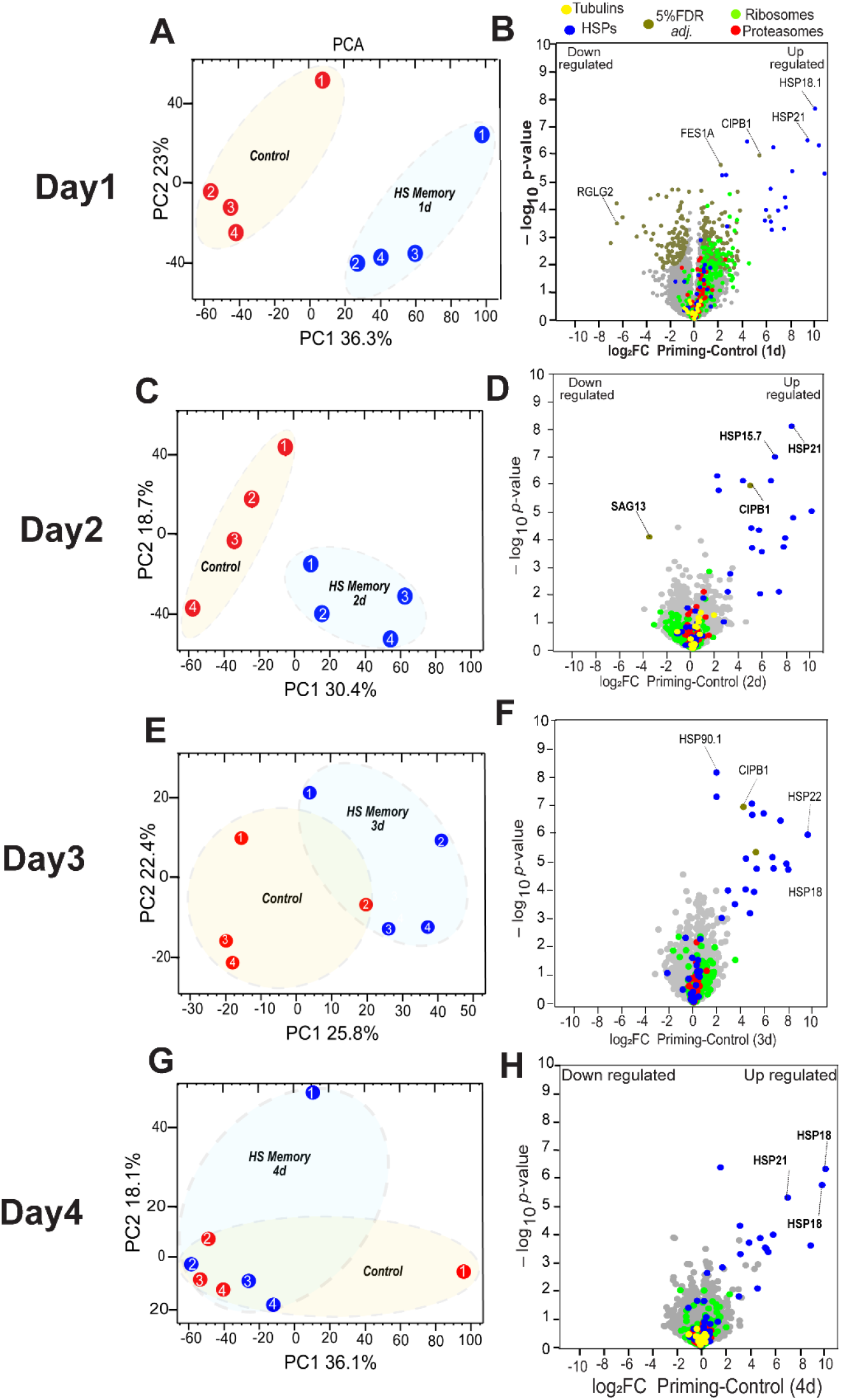
Proteomic analysis of the Arabidopsis seedlings during the HSM phase (A-G) Principal component analysis (PCA) of the proteomic analysis of day1-4 depicting the variances illustrated by each component. The four biological replicates under control conditions and memory phase have clustered well together. Note that clustering of the HS recovery/memory phase and control phase is more evident during the early recovery/memory phase compared to the later memory phase. The red and blue dots indicate control and priming proteins in Arabidopsis. (B-H) The volcano plot displays protein abundance changes in Arabidopsis seedlings between the control and HS memory phase. Each protein’s abundances were plotted based on its log_2_FC (fold change, FC) and its *P*-value (-log_10_) in significance based on four biological replicates. Note that dark blue dots indicate HSPs, dark green dots indicate proteins that passed the 5% false discovery rate, red dots indicate proteasomes, light green dots ribosomes, yellow dots indicate the house keeping proteins (tubulins) and grey dots indicate proteins with no significant change in it.

**Figure 3.**
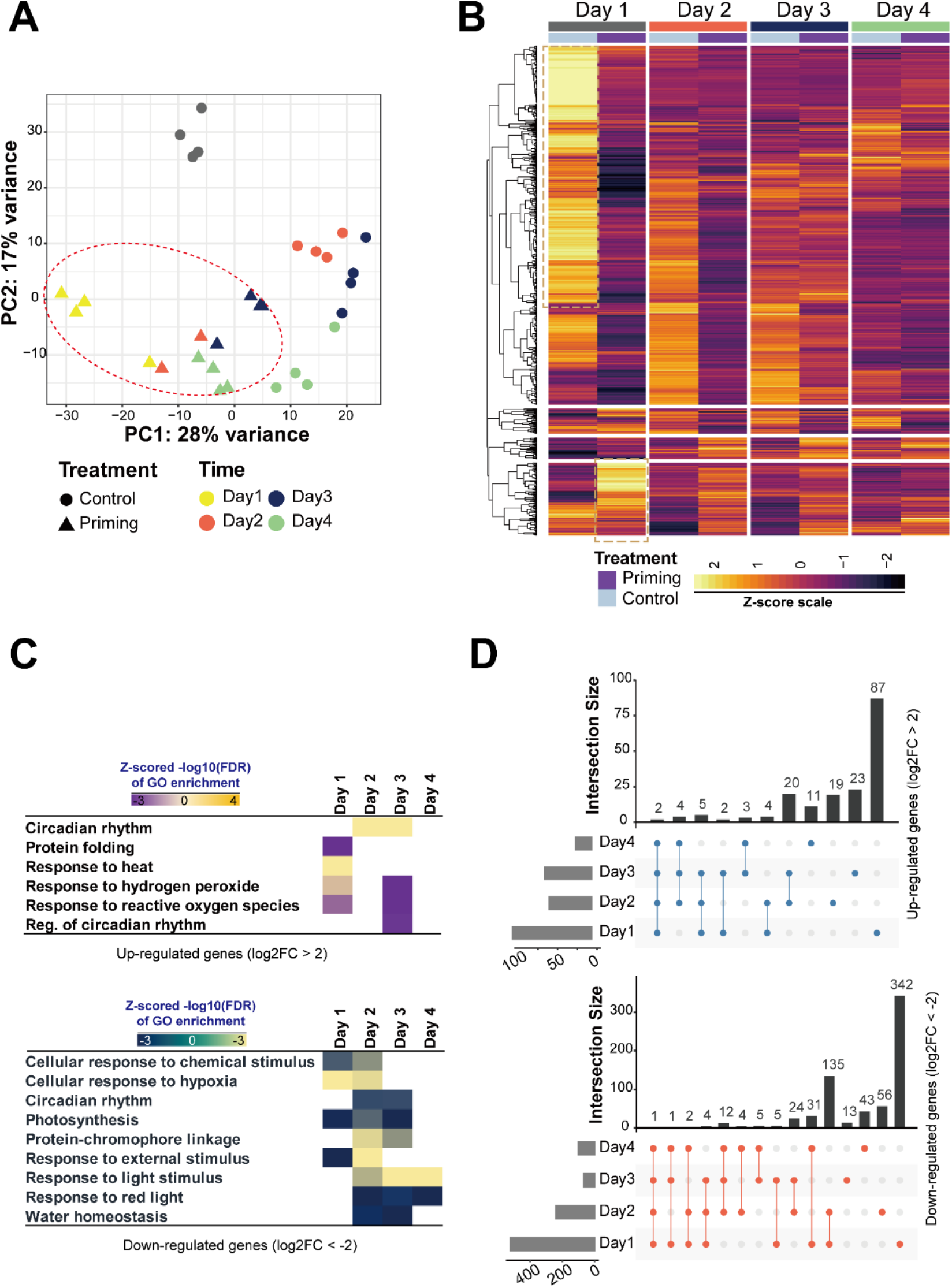
Transcriptomic analysis of HS recovery/memory phase in Arabidopsis. **(A)** Principal component analysis (PCA) of the transcriptomic analysis illustrates the variances by day and treatment. (**B**) Heat map analysis of the differentially regulated genes, clustered by days, scale bar (usually put minus value left and plus value right side) (**C**) Gene ontology (GO) enrichment of the up-regulated genes (upper panel) and down-regulated genes during the HSM phase (lower panel). (**D**) Representation of commonly expressed up-regulated genes within various days of recovery/memory phase. The vertical black bars indicate the number of unique genes at the intersection between different days in the recovery/memory phase (upper panel). Representation of commonly expressed downregulated genes within various days into the HS memory phase. The vertical black bars indicate the number of unique genes at the intersection between different memory days (lower panel).

The differentially regulated transcripts, both upregulated and downregulated are depicted in a heatmap (**Figure 3B**). The gene ontology annotations associated with the upregulated and downregulated genes is shown in (**Figure 3C**, upper and lower panels, respectively). Notably, pathways, including circadian rhythm, protein folding machinery, ROS signaling and response to heat, are enriched in the upregulated categories, while pathways related to photosynthesis, response to hypoxia and protein translation machinery are distinctly presented among the downregulated genes (**Figure 3C**, lower panel). The intersection of the up-regulated and the down-regulated genes was visualized using an upset plot (**Figure 3D**).

Finally, to investigate additional mechanisms and identify novel metabolites associated with HSM, we utilized a metabolite profiling approach. We targeted primary metabolites of Arabidopsis seedlings at different time points during the HS recovery/memory phase alongside their respective controls using GC-MS. Consistent with the transcriptomics and proteomics findings, metabolite changes were more prominent during the initial phase of HSM, with greater impacts on days 1 and 2 compared to days 3 and 4 (**Figure 4A**). Notably, intermediates of the tricarboxylic acid showed upregulation during the HS memory phase (**Figure 4B**). Overall, our multi-omics dataset provides a robust platform for future studies to captivate and understand novel heat stress-related processes.

**Figure 4.**
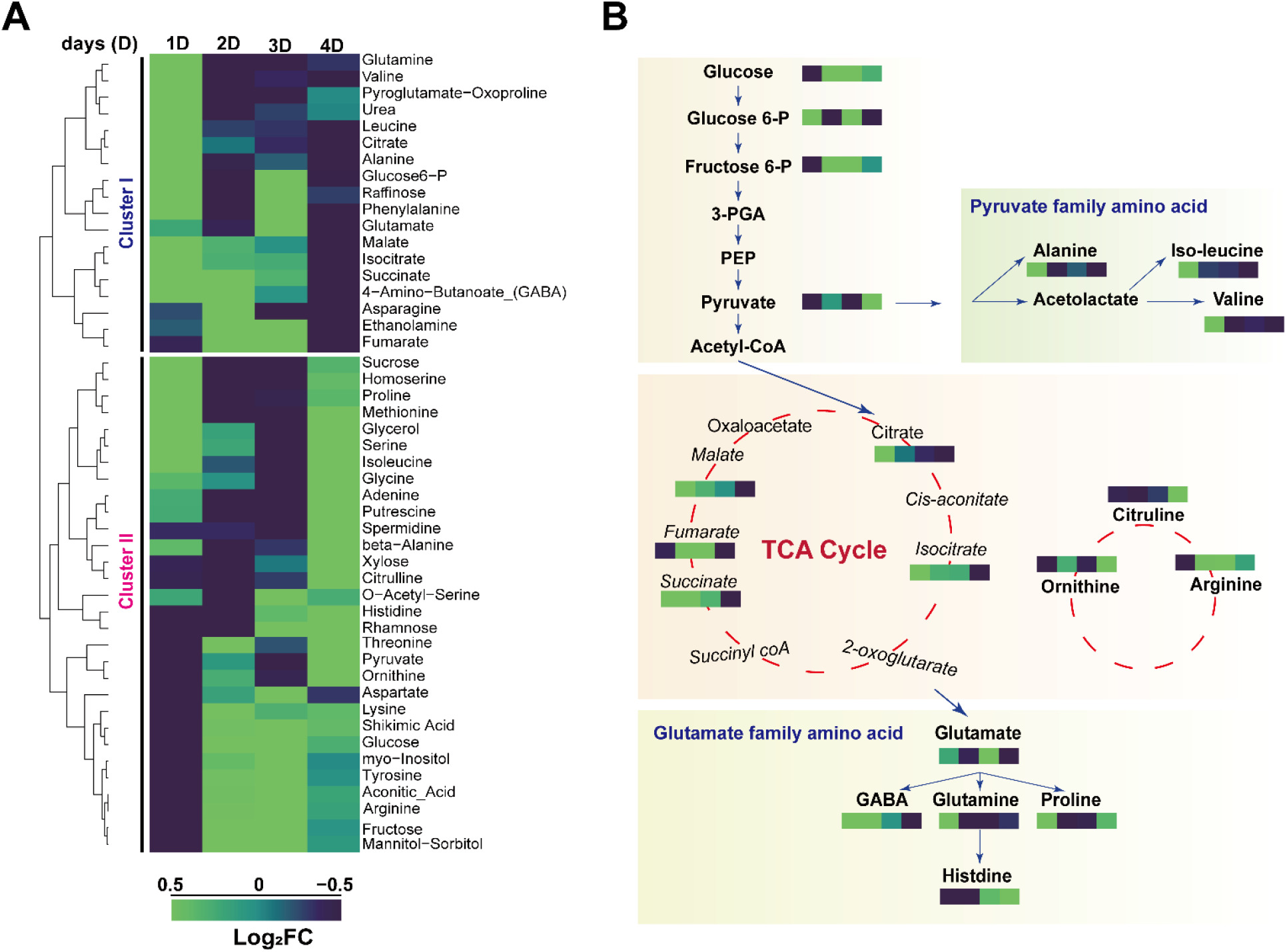
Metabolomic profiling of Arabidopsis seedlings during the Heat stress memory phase. **(A)** Primary metabolite changes in Arabidopsis seedlings are illustrated through a heatmap. The relative abundances of sugars, amino acids, sugars are shown in log_2_ fold change (log_2_fc) scale, with upregulation represented in green and downregulation in violet. Scale bars are as indicated in the figure **(B)** A schematic representation of major primary metabolites from glycolysis to tricarboxylic acid cycle is highlighted in a pathway heat map style. For the detected metabolic features, the relative abundances are depicted.

To visualize the multi-omic relationship between the omics layers across control and priming stimuli, we employed the DIABLO analysis suite. The correlation coefficients between the transcriptomic (mRNA), metabolomic (meta), and proteomic (prot) datasets were visualized using an off-diagonal plot (**Figure 5A**). As expected, we observed a separation between the experimental groups of all pairs, indicating a meaningful integration and alignment among these biological signals. The diagonal panels display the correlation coefficients between components, representing a high level of agreement across datasets (0.88–0.98). We then employed cross-omics to unravel any potential molecular relationships and regulatory mechanisms. As indicated in **Figure 5B**, the circos plot reveals a highly interconnected network of features across the transcriptomic, proteomic, and metabolomic datasets. Interestingly, we observed a higher number of positive correlations (**Figure 5B**, orange links, in the circos plot) in comparison to negative correlations (**Figure 5B**, black links, in the circos plot). Overall, the circos plot indicated notable features from each omic layer correlating with other omic datasets. Next, we observed an unparalleled yet distinct functional relationship between transcriptomic, metabolomic, and proteomic datasets under control and priming conditions (**Figure 5C**), To assess the individual contributions of each omics layer and to better understand the discriminatory power of specific variables driving the separation between the experimental groups, we utilized the variable contribution plot for component 1 and 2 of the DIABLO model. As visualized by the bar plot, the loadings (weights) of selected metabolite features indicate their relative influence on the first latent component that separates the priming and control experimental conditions from each omics layer. The contribution plot for components 1 and 2 shows how individual metabolites, proteins, or transcripts drives the separation between the Priming (orange) and Control (blue) groups (**Figure 5D**). In the metabolomics dataset, contribution 1 plot shows phenylalanine to be influential. Phenylalanine showed the highest positive loading, indicating a strong association with the primed condition. Additionally, the contribution plot for component 2 revealed that pyruvate and lysine were altered under control conditions, suggesting a role for these metabolites in seedling growth and development (**Figure 5D, lower panel**). Similarly, we analyzed the protein-specific loading values to identify key proteins contributing to the separation between primed and control conditions. The most influential protein associated with priming was *AT1G73940*/ML106, followed by *AT5G60250*/C3HC4-type RING finger, *AT3G60090*/VQ26, and *AT1G29120*/UP9, indicating that these proteins are more important in the primed state, as shown in the contribution plot component 1. By contrast, proteins such as Q9FYC4/FAMT were less relevant to priming stimuli, indicating a role in control conditions. For the proteins, at component 2 Q9XIA9/ LACS2 showed the most influential change under control conditions, its level is higher in control than in priming. Interestingly, LACS2 levels are directly linked with Arabidopsis seedling development (Xie et al. 2020) (**Figure 5D, middle panel)**. Finally, at the transcript level, genes such as *AT5G03430*.*1/PAPS, AT5G11560/PNET5*, and *AT5G27030/TPR3* were found upregulated upon priming stimuli, however, *AT1G32190*/Alpha beta hydrolase showed a negative loading (**Figure 5D, upper panel**). Dirigent protein 13 (*At4g11190/DIR13*) was identified as the most influential contributor to component 2, despite the lack of detailed studies on this gene. The potential role of *DIR13* in seedling growth warrants further investigation in future studies. Interestingly, a limited number of transcripts were more prominent in control samples. This supports the observation that priming has a unidirectional effect on the transcriptome, leading to widespread transcriptional reprogramming. This study provides valuable insights into the complex interplay between gene expression, protein synthesis, and metabolic changes during the formation of heat stress memory in plants. The findings have significant implications for developing strategies to enhance crop resilience to increasing temperatures due to climate change.

**Figure 5.**
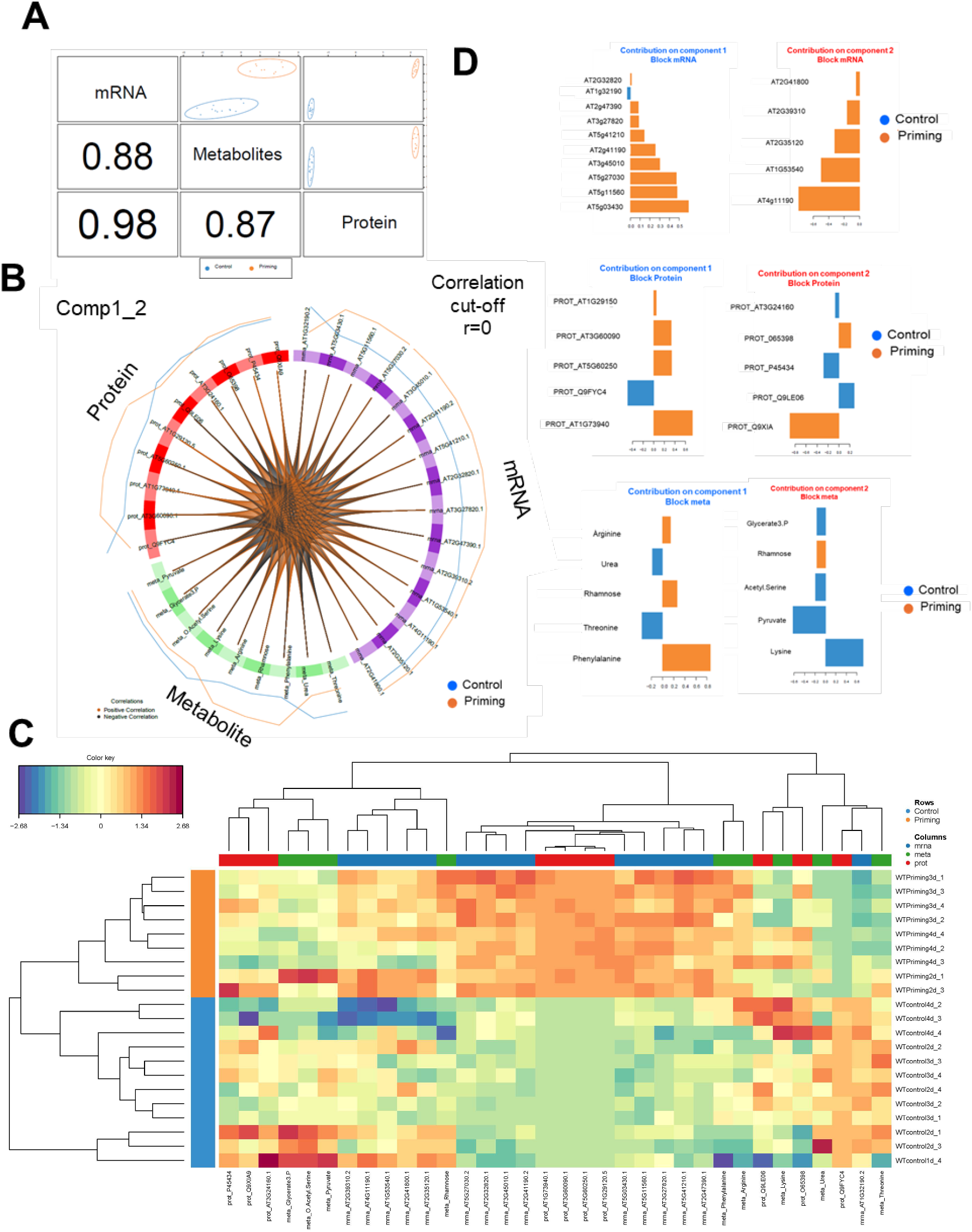
Integration of Multi-Omic Datasets in Arabidopsis During the Heat Stress Memory Phase. **(A)** Offset diagonal plot displaying the correlation coefficients between transcriptomic, proteomic, and metabolomic datasets. The numbers within the panel indicate the correlation coefficients between the respective components. **(B)** Circos plot depicting the functional connections among different omic layers. Orange lines represent positive correlations, while blue lines indicate negative correlations. **(C)** Heatmap illustrating the relationships among the multi-omic datasets. Scale bars are provided in the accompanying color key. **(D)** Contribution plot highlighting the most influential features at the mRNA (top panel), protein (middle panel), and metabolite level (lower panel).

## Experimental procedures

### Plant materials and growth conditions

*Arabidopsis thaliana* ecotype Col-0 seeds were surface sterilized and sown on Petri dishes containing an equal amount of Murashige-Skoog (MS) 0.8% agar medium supplemented with 1% sucrose (w/v). Seeds were stratified at 4°C in darkness for 2 days. Seedlings were grown in growth chambers experiencing a 16 h light/8 h dark cycle at 22°C.

### Heat priming treatment of *Arabidopsis thaliana* seedlings

All priming heat stress treatments reported in the study were performed in Petri dishes as described (Sedaghatmehr et al. 2016; 2019, 2021, Thirumalaikumar et al. 2021). For the priming heat stress, five-day-old seedlings were subjected to a heat regime of 1.5 h at 37 °C (incubator); 1.5 h recovery at 22 °C; and 45 min, 44 °C (hot water bath). After the priming heat stress treatment, seedlings were returned to normal growth conditions for 4 days (termed the HSM phase) during which samples were harvested for analyses.

### Sample collection, total protein extraction and in-solution trypsin digestion for LC-MS/MS analysis

Five-day-old seedlings of Arabidopsis (Col-0) after heat priming treatment particularly during the thermorecovery phase (as indicated in figure 1) and subsequently were ground using liquid nitrogen. Controls without priming treatment were also harvested at the same time points. Phase separation and total protein extraction were done as described by Salem et al. (2016). For protein extraction, solvent-phase separated protein pellet was dissolved in extraction buffer containing (6 M urea, 2 M thiourea, 15 mM DTT, and protease and phosphatase inhibitors) the solubilized proteins were centrifuged at 10,000 g for 5 min and the protein concentration was determined from the collected supernatant by Bradford analysis. Fifty μg of protein extracts were digested in solution using a trypsin/Lys-C mixture (Mass Spec Grade, Promega) according to the manufacturer’s instructions. Digested peptide samples were desalted using C18 stage tips as described (Rappsilber et al.2007).

Peptide mixtures were analyzed by LC-MS/MS using a Q-ExactivePlus high-resolution mass spectrometer connected to an EASY-nLC 1000 system (Thermo-Fisher, Bremen, Germany).

HeLa digests (Pierce 88329) and Cytochrome c digests have been used to monitor the retention time drift and mass accuracy of the LC.

### Label-free quantitative shotgun mass spectrometry and protein identification

Q-Exactive (Thermo Fisher Scientific, Dreieich, Germany) mass spectrometer coupled to an Ultimate 3000 (Thermo Fisher Scientific, Dreieich, Germany) was used for mass spectrometric proteomic analysis. Methods adapted from Fromm et al. (**2016**). In brief, 1-2 µl of sample solution were injected onto a 2 cm, C18, 5 µm, 100 Å reverse phase trapping column (Acclaim PepMap100, Thermo Fisher Scientific, Dreieich, Germany) at a flowrate of 4 µl min^−1^. Peptide separation was done on a 50 cm, C18, 3 µm, 100 Å reverse phase analytical column (Acclaim PepMap100, Thermo Fisher Scientific, Dreieich, Germany). Peptides were eluted by a non-linear 5–30% (v/v) acetonitrile gradient in 0.1% (v/v) formic acid at a flow rate of 300 nl min^−1^ and over a period of 90 min and at 35 °C.

Transfer of eluted peptides into the mass spectrometer was performed using a NSI source (Thermo Fisher Scientific, Dreieich, Germany) equipped with stainless steel nano-bore emitters (Thermo Fisher Scientific, Dreieich, Germany). Spray voltage was set to 2.4 kV, capillary temperature to 275 °C, and S-lens RF level to 50%. The MS was run in positive ion mode, MS/MS spectra (top 10) were recorded from 20 to 100 min. Full MS scans were performed at a resolution of 70,000 whereas 17,500 was used for MS2 scans. Automatic gain control (AGC) targets for MS and MS/MS were set to 1E6 and 1E5, respectively. Only peptides with 2, 3, or 4 positive charges were considered.

### Proteomic data analysis

Raw data were processed using the Proteome Discoverer (version 2.5, ThermoFisher scientific) software using Arabidopsis protein database (Version 10, The Arabidopsis Information Resource, www.Arabidopsis.org). An in-house list of common contaminants was added to the search. SEQUEST HT was used to assign the peptides, allowing a maximum of 2 missed tryptic cleavages. Additionally, a minimum peptide length of 6 AAs, a precursor mass tolerance of 10p□pm and a fragment mass tolerance of 0.02□Da were selected. Carbamidomethylation of cysteines and oxidation of methionine were specified as static and dynamic modifications, respectively. False discovery rates (FDRs) of 0.01 and 0.05 validated peptide spectral matches at high and medium confidence, respectively. Label-free quantification (LFQ) based on MS1 precursor ion intensity was performed in Proteome Discoverer with a minimum Quan value threshold set to 0.0001 for unique peptides. The ‘3 Top N’ peptides were used for area calculation. The normalized protein abundances were used among the samples and values were log2 transformed and imputed using normal distribution pattern. R-studio and Microsoft excel was used to plot the graphs.

### RNA extraction and sequencing

RNA was extracted from *A. thaliana* seedlings during heat stress memory phase and appropriate control conditions as described previously (Balazadeh et al.2008; Thirumalaikumar et al.2021). Briefly, 100 mg of liquid nitrogen ground samples were used to extract total RNA using Trizol reagent followed by Ambion RNA extraction kit (Invitrogen) following manufacturer’s instructions. DNAse I (1u/uL) (Thermofisher) treatments were done following the manufacturer’s protocol. The integrity of the RNA was verified using an RNA gel and the concentration was estimated using Nanodrop.

### Transcriptome sequencing and data processing: (liang)

Four biological replicates of DNASE treated RNA samples during control and heat stress memory phase were sequenced on the HiSeq 4000 platform (Illumina: 100bp single-end). The library preparation and sequencing were performed at BGI Genomics, China (http://www.bgi.com/). Sequencing adaptors and low-quality bases were trimmed using Trimmomatic v0.38 (Bolger et al.2014), and reads shorter than 25 bp were discarded. The filtered short reads were than aligned to the Arabidopsis reference genome using Hisat2 with default settings and the gene level transcription abundance were quantified using FeatureCounts (Liao et al.2014, parameter: -t gene, -g ID, -s 0) based on the latest gene annotation of Arabidopsis (The Araport11, Cheng et al.2017). To ensure the accuracy, we only retained the unique mapped reads to prevent the multiple-counted (located) reads in transcripts. Gene expression profile was performed using the DESeq2 package in R/Bioconductor (Love., et al 2014). Initially, the quantified reads of 61 samples (variable numbers: Treatment (2), genotype (2), timepoints (4), replicates (3∼4)) were normalized using DESeq function. Further, the pairwise comparison of gene expression between the priming and control condition across each timepoint (day1 ∼ day4) were performed to identify differentially expressed genes (DEGs). The criterion of the adjusted P-value cutoff <0.05 and an absolute value of log2 fold change ≥2 were used to classify and filter DEGs. Lastly, the normalized counts were transformed into variance stabilizing transformation (vst) counts, and the top 10% genes along with the most variations across 61 samples were retained to characterize the expression profile by PCA. These RNA-seq raw reads were deposited on NCBI BioProject database (www.ncbi.nlm.nih.gov/bioproject) under ID SUB15359361.

### Metabolite extraction

Primary metabolites were extracted from ground seedling material as described by Giavalisco et al. 2011; Salem et al. 2016. Briefly, approximately 50 mg frozen ground plant material was resuspended in 1 mL of pre-cooled (-20 °C) methyl tert-butyl ether followed by phase separation with 700 μL of UPLC-grade water/methanol (3:1 v/v) and centrifugation. After removal of the organic phase, 200 μL of the polar phase was aliquoted to a fresh 1.5 mL Eppendorf tube and dried in a SpeedVac. Dried samples were derivatized and primary metabolites were then measured by GC-TOF-MS (Caldana et al. 2013). Raw GC-MS files were processed following the pipeline described by Cuadros-Inostroza et al.2009. Further data analysis was computed using Rstudio.

### MultiOmic Integration

To perform supervised integration of multi-omics datasets, we utilized the DIABLO (Data Integration Analysis for Biomarker discovery using Latent components) framework implemented in the mixOmics R package (Singh et al.2019). We collected proteomic (5,861 features), metabolomic (56 features), and transcriptomic (23,253 features) datasets for matched samples. Features with missing values in more than 25% of the samples were removed from each omics layer to ensure data quality. The remaining missing values were imputed using a k-nearest neighbors (KNN) approach. After filtering, the proteomics and metabolomics datasets retained 5,533 and 50 features, respectively, while the transcriptomic dataset retained 20,447 genes. Due to the high dimensionality of transcriptomic data, which included over 23,000 genes, we applied an additional filtering step based on differential expression analysis. Only genes with a false discovery rate (FDR) ≤ 0.05 between priming vs. control groups were retained to reduce noise and contains biological relevance features, resulting in 4,815 transcriptomic features. In contrast, the proteomics and metabolomics datasets contained substantially fewer features (5,861 and 56, respectively), and thus did not require such filtering to ensure model interpretability. Subsequently, each omics dataset was normalized separately using the Trimmed Mean of M-values (TMM) method by calcNormFactors from the edgeR package to account for library size and composition bias Robinson et al.2010 The preprocessed and normalized datasets were then integrated into a multi-block structure and analyzed using DIABLO to identify multi-omics signatures predictive of experimental conditions. The classification performance of the DIABLO model was evaluated using repeated 10-fold cross-validation. Among the three distance metrics tested (maximum, centroid, and Mahalanobis), max.dist consistently showed the lowest classification error rates across both components. The first component accounted for the majority of the discriminative ability, with a balanced error rate below 5%, suggesting a strong and stable separation between classes. Inclusion of a second component provided minimal improvement and slightly increased variability, indicating diminishing returns beyond the first component.

### Data Records

RNA-seq reads and their corresponding peak files have been submitted to Nation center for biotechnology information sequence read archive (NCBI SRA number: SUB15359361) Bio project number: PRJNA1270695. The proteomic raw files have been submitted to the Proteomics identification database massive repository MSV000098061. The metabolomic raw data of the corresponding samples are submitted to massive (MSV000098074).

## Statistical analysis

Statistical analyses were performed using GraphPad Prism (GraphPad Software Inc., San Diego, CA), Experimental data were analyzed with R studio and excel.

## Acknowledgements and Competing Interests

SB thanks the Deutsche Forschungsgemeinschaft (DFG) for funding of the Collaborative Research Centre 973 ‘Priming and Memory of Organismic Responses to Stress’ (www.sfb973.de). Support by the Max-Planck Institute of Molecular Plant Physiology, the University of Potsdam, Cornell University, and Boyce Thompson Institute is gratefully acknowledged. We would like to acknowledge NSF-IOS 2023310 and NSF-IOS 2102120 (to ADLN). The authors used open AI’s ChatGPT (https://chat.openai.com/chat) to adjust the manuscript’s English.

